# Analysis and Design of Frequency-Based Biological Signaling Cascades

**DOI:** 10.64898/2026.08.19.745833

**Authors:** Arya Eimagh Naeini, Shayan Nejad, Devin O’Donnell, Thomas E. Kuhlman

## Abstract

Based on our experimental observation of activation state oscillations of different frequencies used to communicate information by the master human stress response regulator protein p38 MAPK ^1^, we develop a simple graphical approach for understanding and predicting the behavior of complex biological networks acting upon signals carrying information as different frequency waves of chemicals. This approach uses the same techniques used for analyzing and understanding information transmission using waves of electrical currents and fields used in electrical alternating current (AC) circuits. We show how biological components can be organized to behave as standard components found in electronic telecommunications circuits. Finally, we demonstrate how such components can be organized into complex biological signaling cascades whose behavior can be qualitatively and quantitatively understood, and whose output resembles that experimentally observed in p38.

## INTRODUCTION

All living organisms must transmit signals carrying information about the environment to the nucleus to alter gene expression to dynamically respond and adapt. In eukaryotic cells, in many cases this signaling process is accomplished through complex networks of interacting enzymes such as the mitogen activated protein kinase (MAPK) cascade ^2, 3^, the PI3K/AKT/mTOR pathway ^4^, and the Wnt/β-catenin pathway ^5, 6^.

Each of these pathways do not exist in isolation and frequently interact through crosstalk, with, for example, the PI3K/AKT pathway directly inhibiting or activating nodes within the MAPK pathway depending on the stimulus and context ^7–11^. A fundamental question in cell signaling therefore is how core, shared signaling pathways activated by different upstream inputs can produce distinct outputs with unique specificity appropriate to sensed stimuli. While many different mechanisms have been proposed to provide specificity ^12–15^, a growing body of work points to the dynamics of signaling components as playing an important role ^1, 16–24^.

For example, in the MAPK system, an environmental stimulus sensed by a cell starts a complex chain of phosphorylation reactions between MAPK enzymes, adding and removing phosphate residues covalently bonded to specific sites on MAPK components to dynamically alter the enzymes’ activation states ^2, 25–27^. In the case of responding to environmental stresses, we have recently found that the master human stress response protein p38 MAPK encodes information about stimuli as different frequency oscillations of its activation state, and targets are selected through frequency-dependent resonance of oscillating biochemical reactions between p38 and its substrates ^1^. This provides a mechanism for enzyme specificity that is dependent on the dynamical state of the p38 MAPK protein.

Moreover, as the phosphate residues that are added and removed from MAPK components are negatively electrically charged, oscillating MAPK activation states correspond to a literal alternating electrical current flowing through biological enzymes rather than wires. We have shown in the p38 MAPK system how its behavior and substrate selectivity can be described in quantitative detail using exactly the same analytical methods used to model information transmission in different frequency waves of electrical fields and currents in AC circuits ^1^.

Based upon these observations of p38 and thousands of other proteins that have been found to oscillate in response to stimuli ^16, 17, 22, 24, 28–37^, we hypothesize that these complex, interconnected networks of dynamically oscillating proteins generally function exactly as complex AC circuits, encoding and transmitting information about their environment in protein oscillations. Here we develop a graphical language to understand such frequency-based biological signaling networks based on AC circuit analysis methods, where each element of the graph corresponds to a precise mathematical expression describing the impact of that component on the signal. We use this language to show how multimerization and complex formation can generate different frequency oscillatory signals, and we show how to implement a variety of typical AC circuit elements using biological components to filter and isolate different frequency signals. Finally, we show how a complex, interconnected biological network can be constructed by combining these components together to qualitatively and quantitatively understand the overall behavior of the network.

## RESULTS

### Michaelis Menten Binding and Reaction Rates

In the Michaelis-Menten formalism ^38^, the binding of an enzyme, *E*, to a substrate, *S*, to produce a product, *P*, is described by the reaction

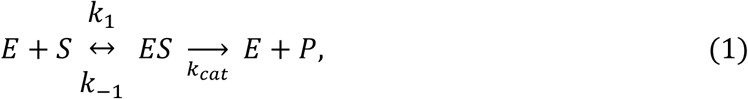

with *k_cat_* the catalytic rate constant and *k*_1_and *k*_−1_ the forward and reverse binding rate constants, respectively. The rate of product formation is

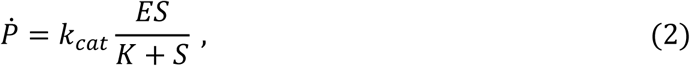

with *E* the enzyme concentration, *S* the substrate concentration, and *K* the Michaelis constant. Dot notation indicates time derivatives. If the substrate is in excess, then *S* >> *K* and the rate only depends on and increases linearly with the enzyme concentration

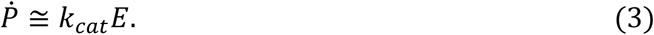

We will assume that all the systems described below are in this linear regime, **eq. 3**.

### Chemical Reactions as a Low Pass Filter

Assuming linear degradation with rate *β*, the effect of a general reaction that does not include any autoregulatory feedback on the rate of production of a component *x* can be written

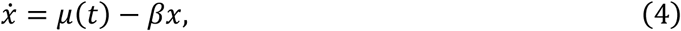

with *μ*(*t*) corresponding to the input from other components affecting the rate of *x* production (**FIGURE 1A**). This component *x* could be protein, RNA, or any other biological element.

**Fig. 1.**
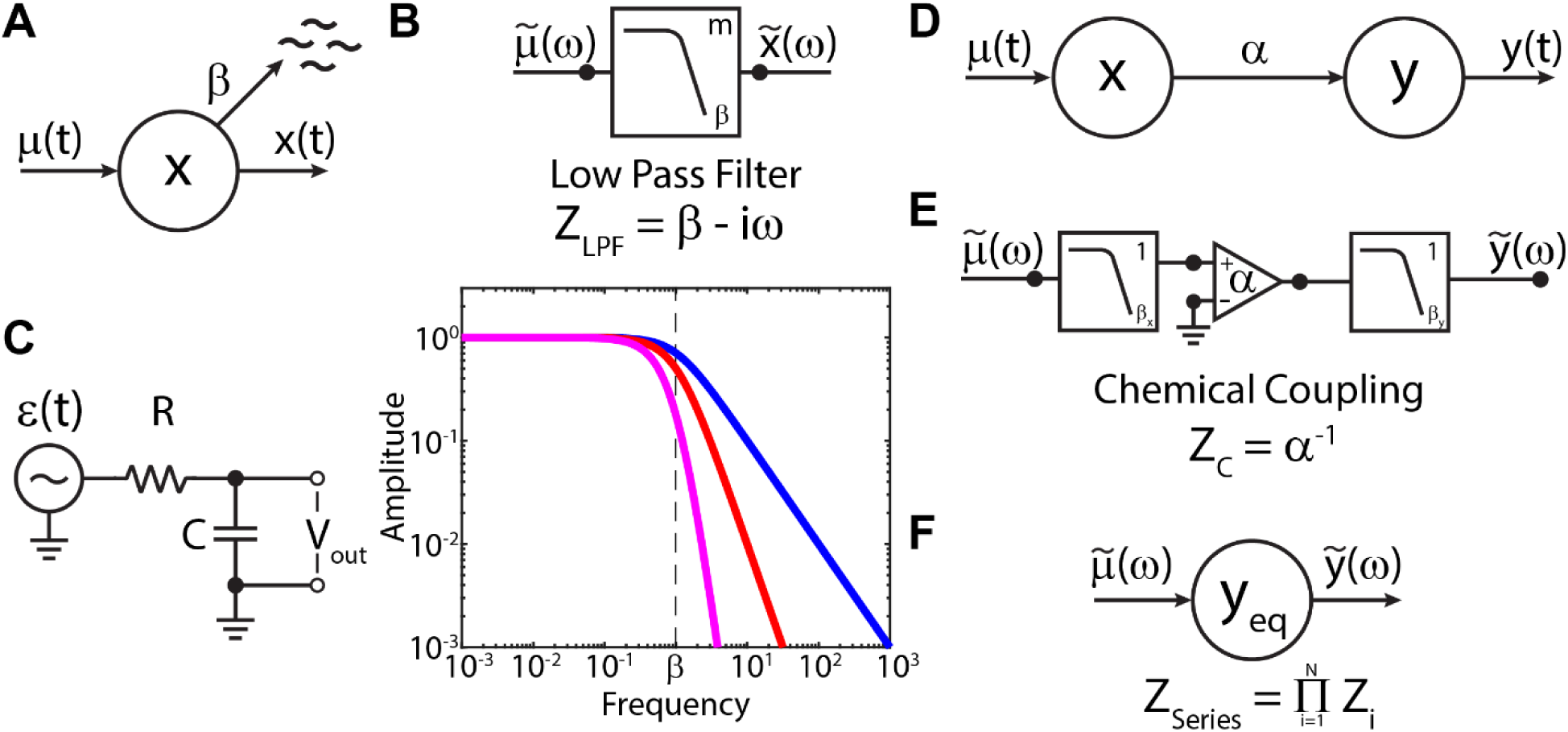
Coupled general chemical reactions as low pass filters coupled by op-amps. (A) A general chemical reaction of component *x* that degrades with linear rate *β*. Input to the reaction is represented by *μ*(*t*) and the output is the time dependent concentration of *x*, *x*(*t*). (B) Top: the effect of this reaction on an input signal corresponds to a first order (*m* = 1) low pass filter. Bottom: amplitude of the output of *x* as a function of input frequency with *m* = 1 (blue), *m* = 2 (red) and *m* = 5 (magenta). (**C**) Equivalent electronic low pass filter implemented with an RC circuit. The output is taken as the voltage across the capacitor, *C*. (**D**) Two coupled chemical reactions, where component *x* is used to drive component *y* with rate constant *α*. Linear degradation of each component will subsequently be assumed and not graphically represented. (**E**) Equivalent representation of (D), with the coupling between the reactions represented as an operational amplifier (op-amp, triangle symbol). (**F**) The three components in series in (E) can be combined into a single equivalent element with impedance equal to the product of the individual impedances in series.

It is possible to analyze such components to determine their frequency-dependent behavior with either Fourier or Laplace transforms. The Fourier transform describes the steady state, forced response of the system, while the more general Laplace transform also describes transient, non-steady state elements of the response and depends on initial conditions. We have found experimentally with the p38 MAPK system that the cell encodes information about received stimuli in the steady-state oscillatory frequencies of components ^1^. Consequently, here we will assume that other elements behave similarly, and we will therefore use the simpler Fourier analysis of components’ behavior. Generalization to the Laplace formalism is straightforward by making the substitution −*iω* → *s* = *a* − *iω* throughout.

Taking the Fourier transform and solving for the frequency spectrum of *x*, *x̃*(*ω*), in terms of the spectrum of the input, μ̃(*ω*),

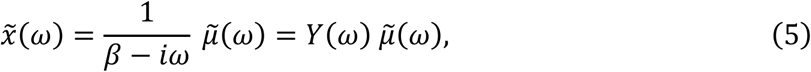

where *ω* is the angular frequency of the signal, *ω* = 2*πf*. This defines the transfer function, *Y*(*ω*), which is a complex function describing how the spectrum of the output arises as a result of the spectrum of the input ^1^. *Y*(*ω*) describes both the amplitude of the response as well as the phase. In many cases the primary interest will be the amplitude of the signal passing through an element, with the phase being only a secondary concern. We see that by expressing *Y*(*ω*) in polar form,

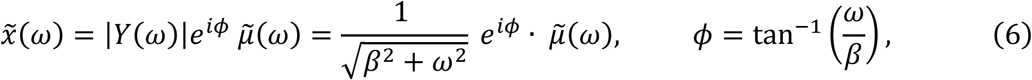

we need only to calculate the magnitude of the transfer function if we are primarily interested in the amplitude of the response.

The amplitude response of this general chemical reaction corresponds to what is referred to in electronics as a first order low pass filter ^39^. The linear degradation rate *β* acts as a cutoff frequency; signals with frequencies below *β* are passed through the component, while those with frequencies above *β* are weakened with amplitude decreasing as ∼*ω*^−1^(**FIGURE 1B**). An exactly equivalent electronic low pass filter can be realized with an RC circuit containing a resistor (R) and capacitor (C) ^40^ (**FIGURE 1C**). The voltage input to the RC filter is *ε*(*t*), and the output of the filter is taken as the voltage across the capacitor, *V_out_*. From Kirchoff’s voltage loop rule, we obtain

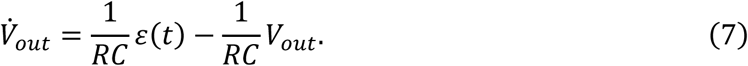

Note the similarity to **eq. 4** above, with *x*(*t*) = *V_out_*(*t*), *β* = 1/*RC*, and *μ*(*t*) = *βε*(*t*). Taking the Fourier transform and solving for the output spectrum ỹ*_out_*(*ω*),

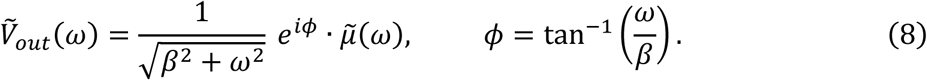

Thus, the transfer function of an RC electronic first order low pass filter is identical to **eq. 6** and results in the same effects on transmission of an oscillatory signal.

In electronic circuit analysis, the transfer function, *Y*(*ω*), is commonly referred to as the admittance, and the inverse of the admittance the impedance, *Z*(*ω*) = *Y*^−1^(*ω*). Hence for a general chemical reaction with linear degradation, **eq. 4**, the impedance is

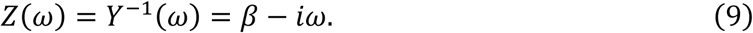

Consequently, **eq. 5** can be written in terms of the impedance as

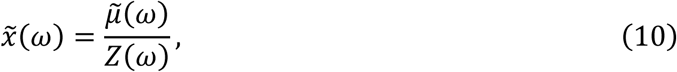

or the AC equivalent of Ohm’s law, with μ̃(*ω*) the input spectrum and *x̃*(*ω*) the output spectrum. Hence, to understand the input-output relationship of a complex electrical or biochemical pathway, it is only necessary to determine the overall impedance of the pathway, which in turn is determined by the impedance of each element along the path.

### Reaction Coupling as an Operational Amplifier (Op-Amp)

Now consider a component *x* with input *μ*(*t*), exactly as above. We take the output of this reaction, *x*(*t*), and use it to drive with rate constant *α* a second component, *y*, which also decays linearly with rate *β_y_* (**FIGURE 1D**). We have

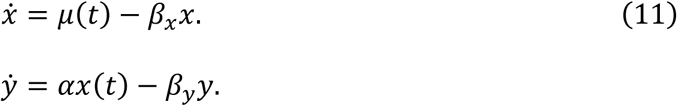

As described above, *x*(*t*) is analogous to the output voltage of an electronic low pass filter. Hence, the input to the second stage component *y* is the previous stage’s output voltage, *x*, multiplied, or amplified, by the rate constant *α*. This coupling between chemical reactions corresponds electronically to the behavior of an operational amplifier, or op-amp (**FIGURE 1E**). A simple op-amp takes two inputs, a non-inverting input, *V*_+_, and an inverting input, *V*_−_, and multiplies the difference of the inputs by the open loop gain, *A_OL_*. The output of the op-amp is

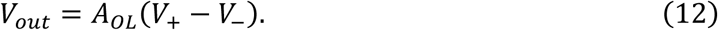

Call the open loop gain *α*. If one of the inputs is taken to be ground, *V* = 0, and the other is taken as an input signal, *x*, to either of the input terminals of the op-amp, then the possible outcomes are

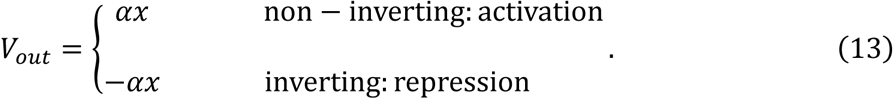

Coupling to a non-inverting output therefore corresponds to the component *x* enhancing or activating the rate of production of component *y*, while coupling to an inverting output corresponds to component *x* repressing component *y*.

The input-output relationship of the coupling between two chemical components corresponds to

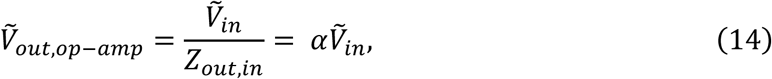

and the impedance of the coupling is therefore *Z*(*ω*) = *α*^−1^.

### Linearly Coupled Reactions and Components in Series

The solution to the coupled *x* and *y* chemical reactions in **eqs. 11** is found by taking Fourier transforms and solving for the output spectrum, ỹ(*ω*),

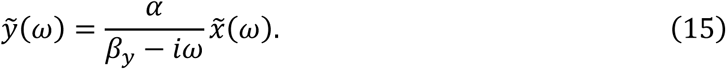

The input spectrum to this reaction, *x̃*(*ω*), can itself be written in terms of its input, μ̃(*ω*), resulting in

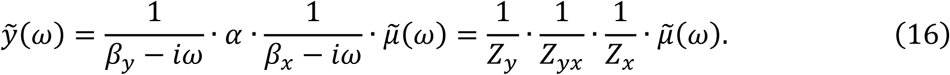

That is, the net effect of these coupled chemical reactions is to divide the original input signal by the product of the impedances of each element. These components directly connected to one another in a linear chain are said to be in series.

The above argument can be iterated for any number of elements. Thus, for any chain of *N* components reacting in series, the final output, *x̃_out_*, can be expressed in terms of the original input, μ̃(*ω*), as

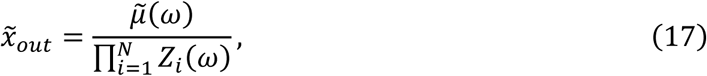

where *Z_i_*(*ω*) are the impedances of each component along the chain. Another way to view this result is that any number *N* of components connected in series can be replaced by a single component with equivalent impedance, *Z_Series_*(*ω*) (**FIGURE 1F**),

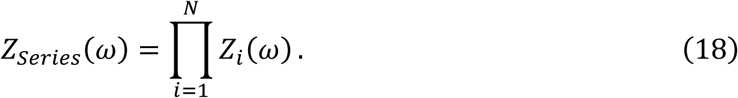

From this, we also see that chains of reacting components act as higher order low pass filters, where the amplitude of the response after passing the frequency cutoff falls off faster than *ω*^−1^. For example, consider an input *μ*(*t*) into a linear chain of *m* identical components with the same decay rate *β* and rate constant *α*. The *n*^th^ component will obey *ẋ_n_* = *αx_n_*_−1_ − *βx_n_*, and the output of the last element in the chain will yield

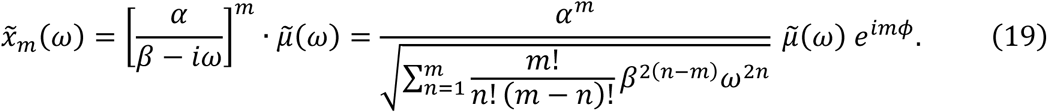

This results in the amplitude dropping off rapidly as ∼ *ω*^−*m*^ past the frequency cutoff *β*, corresponding to an *m*^th^ order low pass filter (**FIGURE 1B**).

### Branched Reactions and Components in Parallel

Consider a single input *μ*(*t*) driving two components, *x* and *y*, which then both independently drive a third component *z*. This biological network is shown in **FIGURE 2A**, and its equivalent representation in **FIGURE 2B**. Because *x* and *y* are not directly linked and are only connected by feeding into the third component *z*, *x* and *y* are said to be connected in parallel.

**Fig. 2.**
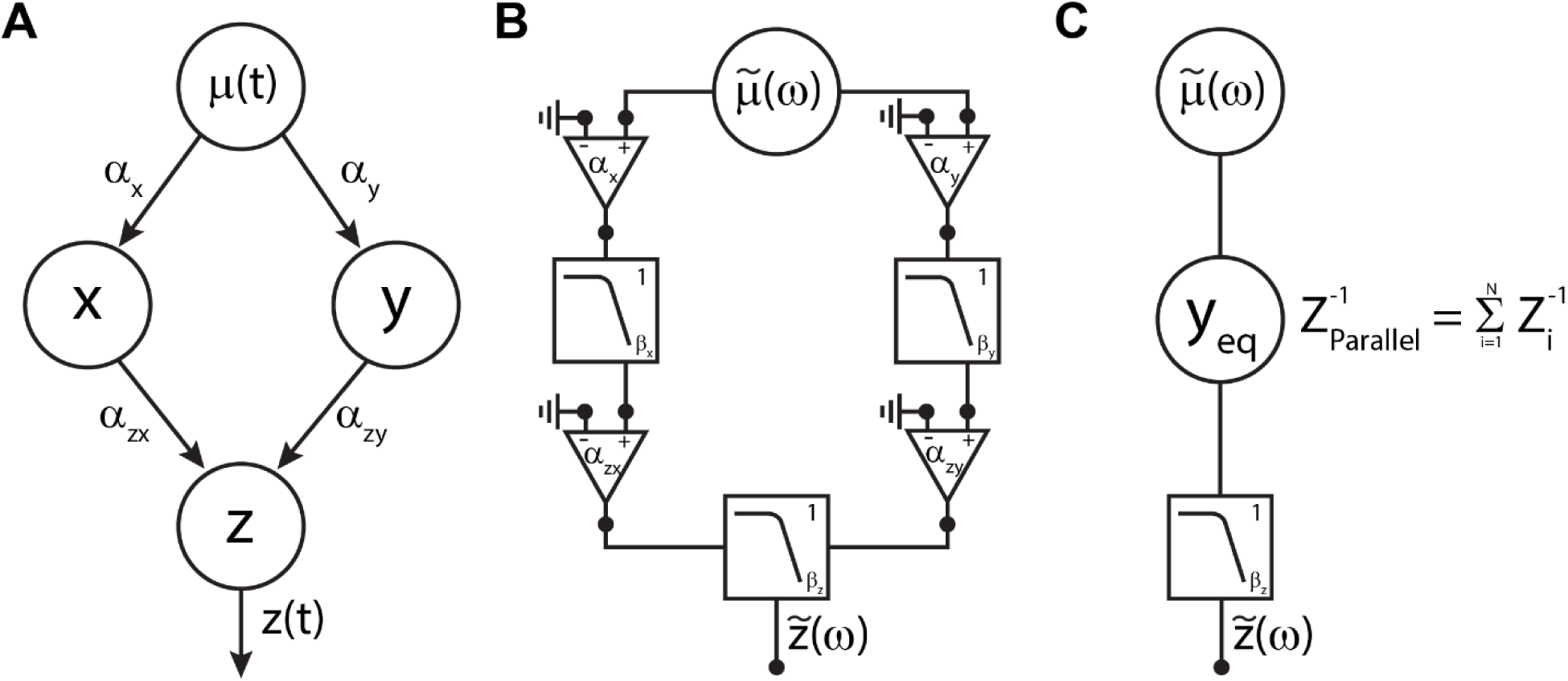
Branched chemical reactions as components in parallel. (**A**) A single input, *μ*(*t*), is used to drive two independent components, *x* and *y*, each of which subsequently drives a third component *z*. (**B**) Equivalent representation of (A). (**C**) The two parallel branches in (B) can be replaced by a single equivalent component whose reciprocal impedance is the sum of the individual reciprocal impedances of each branch.

Each of the parallel branches independently drive *z*, resulting in

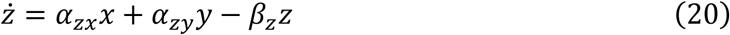

The solution in frequency space is then

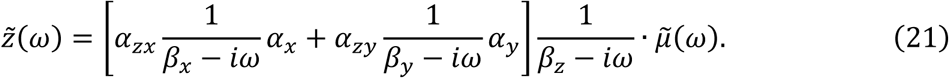

Each of the terms in brackets represents the equivalent impedances of the components in series along each parallel branch, *Z_Xeq_* and *Z_Yeq_*, **eq. 18**. Hence,

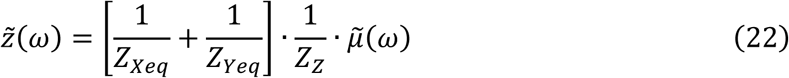

This argument can be repeated iteratively to conclude that *N* branches in parallel can be replaced with a single equivalent impedance as (**FIGURE 2C**)

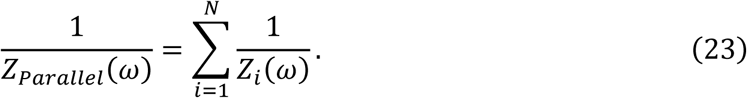

### Biological Implementation of Band Pass and High Pass Filters

In addition to low pass filters, other types of elements frequently employed in AC electronic circuits are band pass filters, where only signals in a frequency range between a low and a high frequency cutoff are passed through the filter, and high pass filters, where only signals with frequencies above a cutoff frequency are passed through the filter.

A biological band pass filter can be implemented using two components, *x* and *y*. Each of these, with linear degradation rates *β_y_* > *β_x_*, are driven by an input *μ*(*t*) that oscillates with frequency *ω*. *x* then represses *y*, and the output of the system is taken as *y*. This is presented diagrammatically in **FIGURE 3A**, with its equivalent representation in **FIGURE 3B**. Note that the couplings of the input to *y* alone and through *x* are two branches in parallel, and that the *x* branch includes multiple components in series. Using the formalism developed above we can immediately read off the resulting overall equivalent impedance

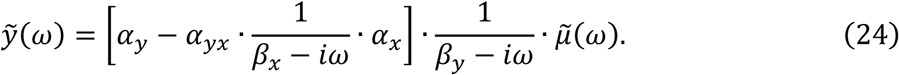

**Fig. 3.**
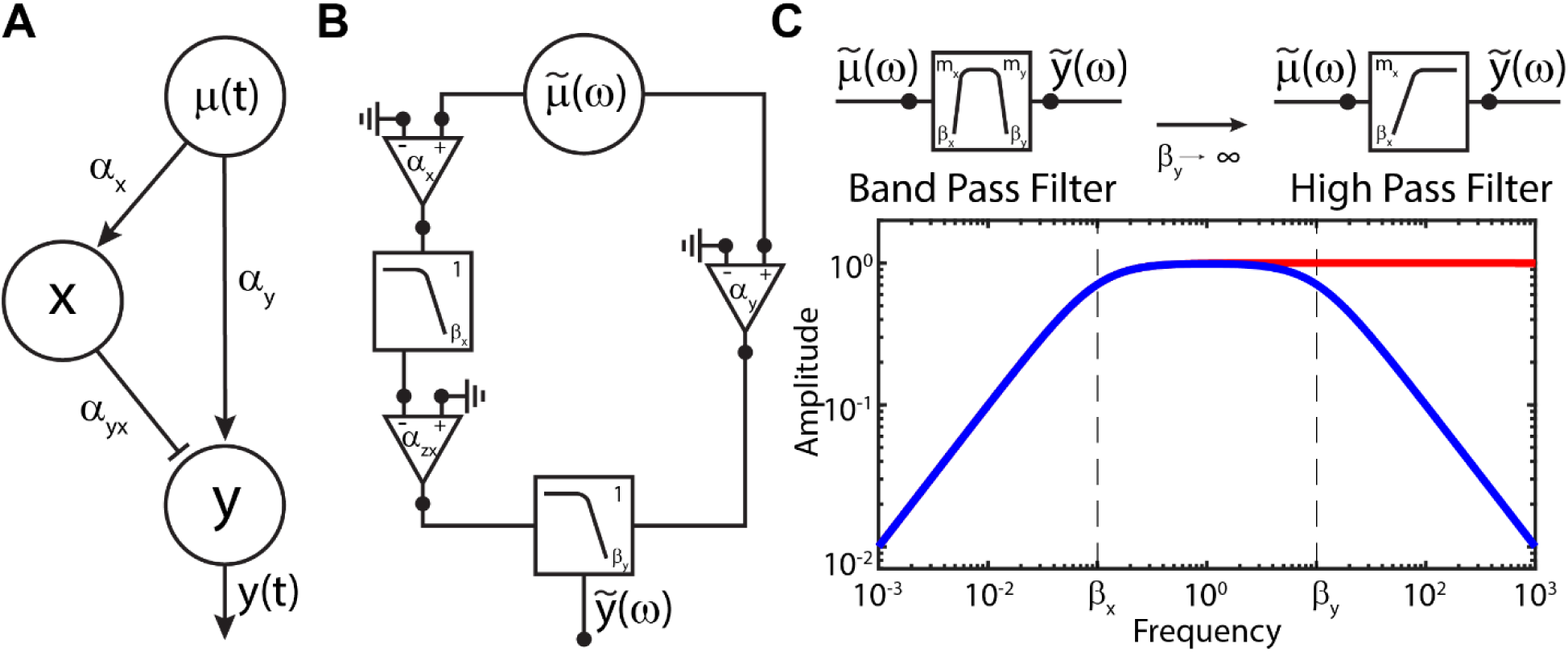
Biological implementation of band pass and high pass filters. (**a**) A single input, *μ*(*t*), is used to drive two independent components, *x* and *y*, and *x* is used to repress *y*. (**B**) Equivalent representation of (A). The output of this system, *y*(*t*), corresponds to a band pass (left, *β_x_* < *β_y_* < ∞) or high pass filter (right, *β_y_* → ∞). Bottom: amplitude of the output signal, *y*(*t*), for a band pass filter (blue) and high pass filter (red).

A plot of the amplitude of this transfer function is shown in **FIGURE 3C**, with *β_x_* < *ω* < *β_y_* the bandwidth through which frequencies pass unattenuated. An intuitive interpretation is that *x* acts as a low pass filter, such that only low frequency components of *μ*(*t*) appear in *x*. These low frequencies are then suppressed in *y* through the repressive action of *x*. But *y* itself also acts as a low pass filter with frequency cutoff *β_y_*. Therefore, if *β_y_* > *β_x_*, there is a frequency band through which signals pass. We also note here that when integrating signals from multiple channels into a single receiver component, it is necessary to consider their relative phase. For example, the effect of *x* suppressing signals in *y* can be achieved either through an inverting input where the two signals are in phase, *i.e.*, *φ_x_* = tan^−1^(*ω*⁄*β_x_*) = 2*πn*, *n* = 0, 1, 2, …, or equivalently coupling through a non-inverting input where the two signals are exactly out of phase, *i.e.*, *φ_x_* = tan^−1^(*ω*⁄*β_x_*) = (2*n* + 1)*π*/2, *n* = 0, 1, 2, … .

A high pass filter, where only frequencies higher than a cutoff are allowed through, can be implemented by *β_y_* → ∞ (**FIGURE 3C**). A band stop filter, where frequencies pass except between a window *β_x_* < *ω* < *β_y_* can be implemented using a third component, *z*, also driven by the same input *μ*(*t*), and coupled to *y* through an inverting, repressive input.

### Feedback Loops and Resonant Amplifiers

Consider the typical two-component biological oscillator model shown in **FIGURE 4A** ^16, 17^. Component *x* receives an input *μ*(*t*) and then positively drives component *y*. Component *y* then represses component *x*. We take *x* as the output. Here, for simplicity, we absorb the coupling rate between the input and *x* into the definition of *μ*(*t*).

**Fig. 4.**
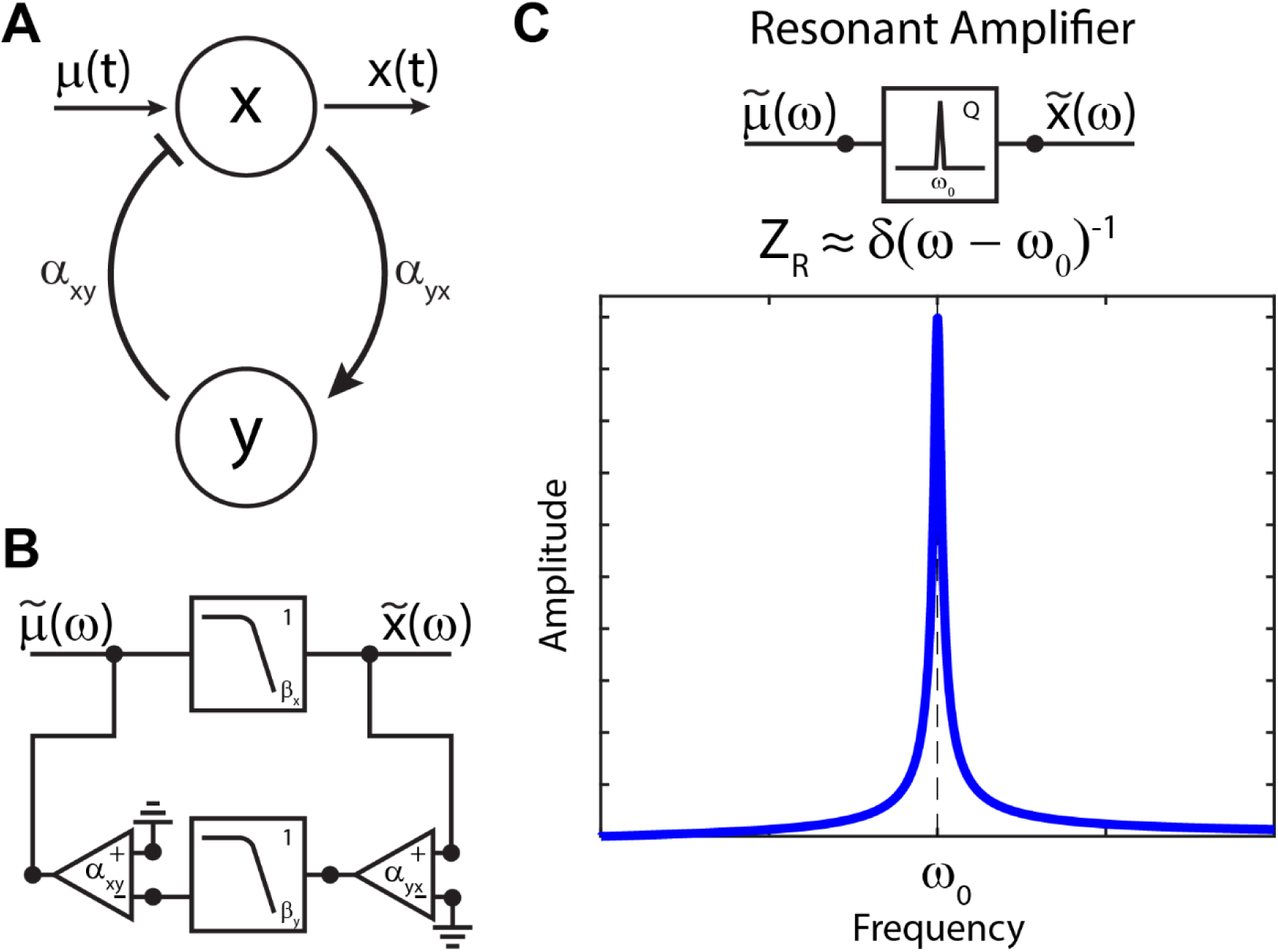
Feedback loops and resonant amplifiers. (**A**) A two-component biological oscillator model, with input *μ*(*t*) to component *x*. *x* acts positively on component *y*, which then feeds back negatively into *x*. *x*(*t*) is taken as the output. (**B**) Equivalent representation of (A). (**C**) A two-component biological oscillator functions as a resonant amplifier, amplifying frequencies only very close to the resonant frequency of the system. Bottom: amplitude of the signal as a function of the input frequency, with *ω*_0_ the resonant frequency.

The equivalent representation of this biological oscillator is shown in **FIGURE 4B**. The branch including *y* constitutes a feedback loop. From control theory ^41^, if the branch containing *x* has impedance *Z_x_*(*ω*) and the feedback branch has impedance *Z_y_*(*ω*), then the overall transfer function describing the output *x* is

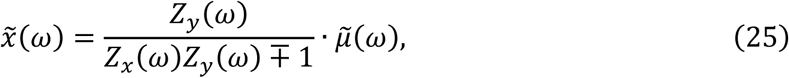

where the minus sign is used if the feedback of *y* into *x* is non-inverting (activating) and the plus sign if it is inverting (repressing).

Complex feedback loops can be progressively simplified using the equivalent circuit procedures outlined above. Here, the main branch includes only component *x* with impedance *Z_x_*(*ω*) = (*β_x_* − *iω*). The feedback branch includes two couplings and component *y* in series, which can be reduced to the equivalent impedance *Z_y_*(*ω*) = *α_xy_*^−1^ · (*β_y_* − *iω*) · *α_yx_*^−1^. Substituting these expressions into **eq. 25** and simplifying yields the impedance of the entire system

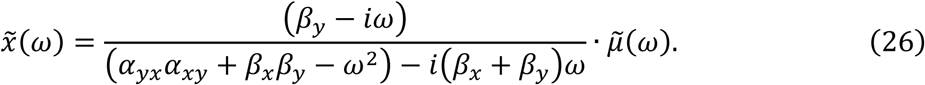

We identify 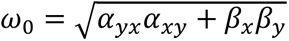 as the natural frequency and *γ* = *β_x_* + *β_y_* as the damping parameter of a driven, damped oscillator system ^1^. The transfer function becomes

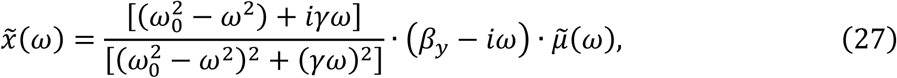

which is recognized as the standard transfer function of a resonant amplifier whose amplitude response is shown in **FIGURE 4C** ^1^.

This system will select and amplify frequencies near the resonant frequency of the system, *ω_r_*, found as the oscillatory parts of the poles of the transfer function

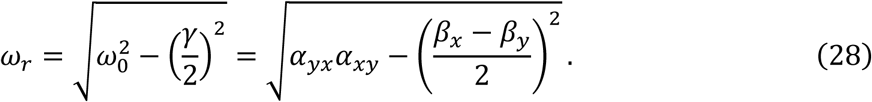

If damping is weak, *β_x_*, *β_y_*, *γ* ∼ 0, and 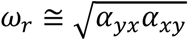

The full width at half max of the resonance peak, Δ*ω*, determines the range of frequencies amplified by the resonant amplifier. The width and height of this peak is quantified by the Quality factor, *Q*,

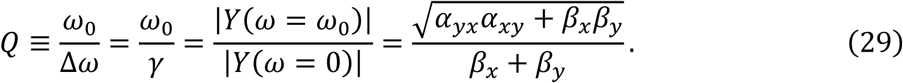

Consequently, if *Q* is large, corresponding to low linear degradation rates, the resonant amplifier will select and amplify only a very narrow range of frequencies centered around the resonant frequency. In the limit of vanishing damping, *Q* goes to infinity (“high-*Q*”) and the response becomes a Dirac delta function spike, *Y*(*ω*) = *Z*^−1^(*ω*) = *δ*(*ω* − *ω_r_*), selecting and amplifying only exactly the natural resonant frequency, *ω_r_*.

### Receptor Binding, Pathway Activation, and Signal Synchronization

To activate signaling pathways, signals received from the environment are typically first detected by membrane-bound receptor proteins. This will result in an input to the signaling pathway of white noise from the stochastic binding and unbinding of the signaling ligands, yielding a constant power spectrum *g̃_input_*(*ω*) = *C*. There are then many possible ways to subsequently organize a network to act upon this noisy input to generate specific frequency oscillatory signals. We hypothesize that oscillators acting as resonant amplifiers should appear early in the cascade to generate specific frequency signals from the noisy input. We will assume, like the p38 system ^1^, that any oscillators are linear and high-*Q,* and hence generate approximately delta function signals in the spectrum at their resonant frequencies. Subsequent layers of the network can then act upon those oscillations to shift, filter, and sculpt the spectrum.

A significant concern is that, if multiple receptors are simultaneously bound and activated, each will create its own stochastic input to a local pool of linear high-*Q* oscillators due to random binding and unbinding of ligands. This will result in a collection of unsynchronized oscillatory signals with random phases. We show here that any form of mixing among these populations of oscillators with random phase, either through passive diffusion or active transport, will result in their synchronization into a coherent oscillatory signal at a single frequency.

We will build the argument in successively more complex steps. First, consider a single receptor activating a local pool of two-component oscillators by giving the *x* component a single random impulse “kick,” instantaneously injecting an amount Δ*x* of active *x*. The mean behavior of the pool after the kick can be written

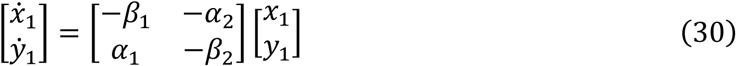

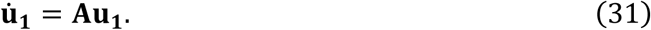

This has the solution

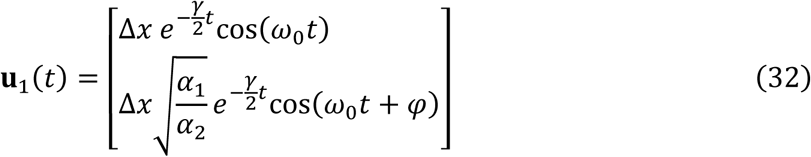

with *γ* = *β*_1_ + *β*_2_, 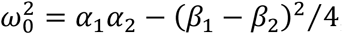 and *φ* = tan^−1^[2*ω*_0_⁄(*β*_1_ − *β*_2_)].

Next, imagine there exists a second receptor similarly providing a random kick to another pool of identical oscillators. Then the second pool will behave identically to the first with solution **u**_2_(*t*), but with a different, random phase.

Now, assume that the pools mix, characterized by a diffusion constant *D*. Then

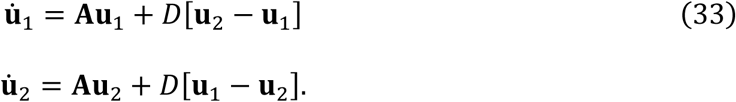

Consider the mean and difference of the two pools,

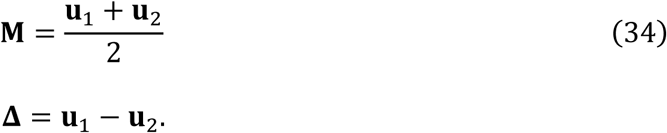

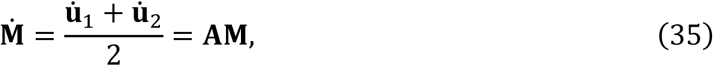

that is, the mean population oscillates identically to a single pool. Conversely, the difference obeys

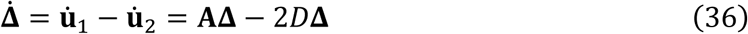

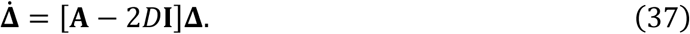

This demonstrates that the difference between the two populations is governed by the same Jacobian as the mean population, but with additional damping due to the mixing, *i.e.*,

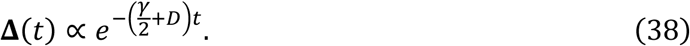

If the diffusion constant *D* is large, the difference between the populations will decay much faster than the amplitude of the mean population’s oscillations, and hence the two pools will synchronize.

Now, rather than a single initiating kick, consider that random activating kicks arrive with average Poissonian rate *λ*. This effect can be expressed as a chemical Langevin equation with an input noise *η*_1_ representing the random input to the *x* oscillator from the receptor. We will assume that the noise is uncorrelated Gaussian white noise,

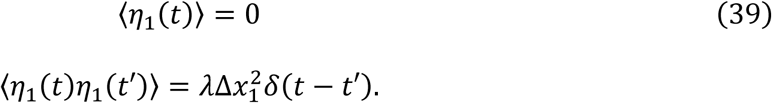

Then

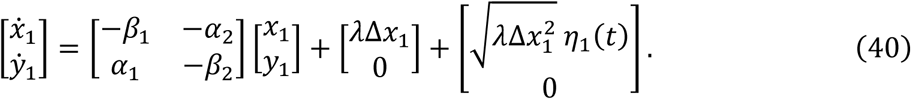

We proceed with the same argument, calculating the mean and difference of the populations. Since the two pools are independent, the noise amplitudes add, yielding for the difference

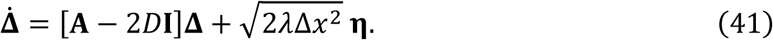

Because of the noise, we expect that the two pools will imperfectly synchronize with some non-zero variance. To quantify this expectation, we define a covariance matrix, **Σ**, and a diffusion matrix, **Q**,

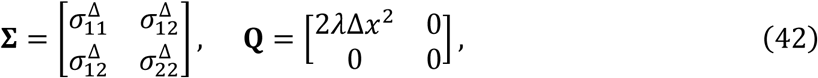

these matrices satisfy a differential Lyapunov equation ^42^,

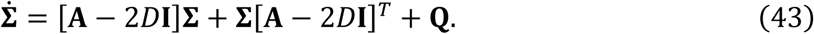

We expect the covariance will reach a steady state,

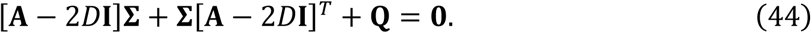

The solution of this Sylvester equation is straightforward. Defining *γ_i_* = *β_i_* + 2*D*, we obtain

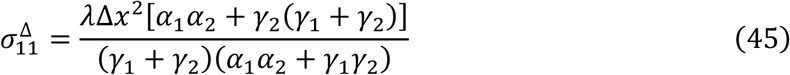

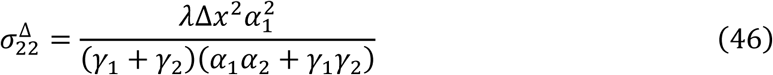

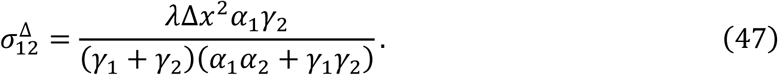

Hence, if the mixing rate *D* is large, these variances will approach zero, corresponding to synchronization.

Finally, we generalize this argument to *N* receptors, each randomly activating *N* local pools. We assume that all pools can mix and combine characterized by the same diffusion constant *D*. As before, the rate of change of each pool must obey

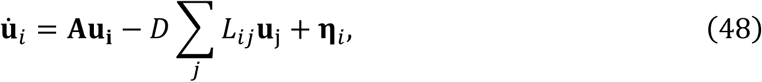

where, for a mean field model where all pools are connected,

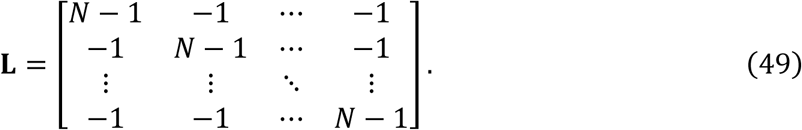

The output of this system will be the mean field

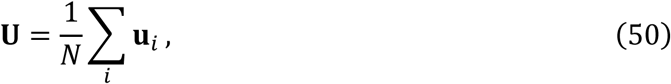

along with the difference modes

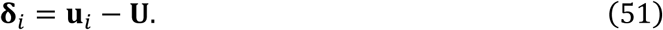

The behavior of the mean field can be found by summing all the individual uϗ*_i_* and dividing by *N*,

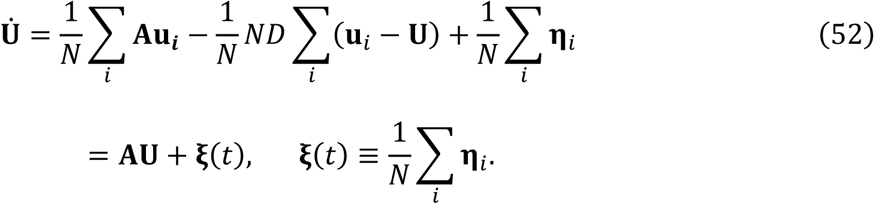

That is, the mean field acts like an oscillator with noise, but the overall noise is divided by a factor of *N* relative to the noise in an individual pool. Repeating the argument to calculate the variance, we obtain

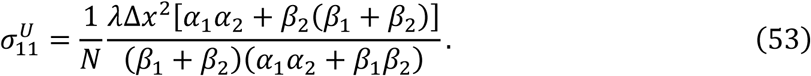

that is, the variance in the mean field due to the oscillations is also reduced by *N*. Consequently, *N* cannot get too large or the average amplitude of the oscillations will approach zero.

For the variance of the difference modes we obtain

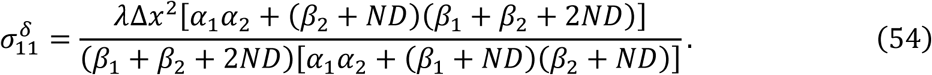

For example, with many receptors and fast diffusion,

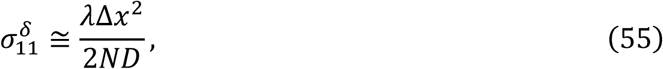

which approaches zero as *N* and *D* get large, corresponding to synchronization.

From this analysis, we conclude that, if there is a large (but not too large) population of receptors, and if oscillators randomly activated by those receptors are mixed rapidly, either through diffusion or active transport, then the mean field output of this system will be a single, coherent oscillatory signal with the same frequency as the individual oscillators.

### Multimerization and Complex Formation as a Biological Heterodyne

Many biological systems exhibit oscillations with periods of many hours, such as the p53 system ^16^, the p38 system ^1, 24^, and circadian oscillations ^43–45^. How do such long periods arise from the high frequency chemical interactions of molecules? The same frequency mismatch problem arises in electronic telecommunications. For example, in FM radio transmissions, low frequency audio information (∼10^1^ - 10^4^ Hz) must be encoded in much higher frequency radio waves (∼10^8^ Hz).

Shifting between otherwise incompatible frequency ranges in telecommunications systems is accomplished through a process called heterodyning. A basic heterodyne consists of three elements: (1) two high frequency oscillators with similar but unequal frequencies *ω*_1_ and *ω*_2_; (2) a nonlinear frequency mixer that combines the high frequency oscillator signals to create new signals at the high frequency sum, *ω*_+_ = |*ω*_1_ + *ω*_2_|, and low frequency difference *ω*_−_ = |*ω*_1_ − *ω*_2_|; and (3) a filter to isolate the desired frequency component.

Consider protein monomers *x* and *y*, whose active levels sinusoidally oscillate at some angular frequencies *ω_x_* and *ω_y_*. Then

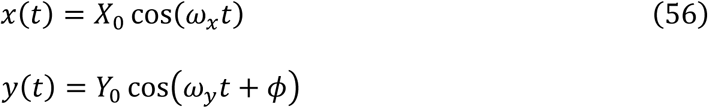

with *φ* an arbitrary phase offset between the two. For simplicity we set *φ* = 0 without substantive loss from the argument. Now suppose these monomers dimerize with rate constant *α* to form a functional complex, *C*, which can then spontaneously disassemble with linear rate constant *β* (**FIGURE 5A**). Then

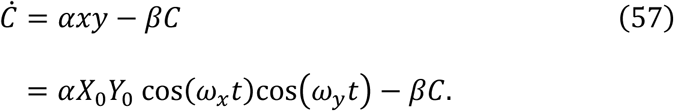

**Fig. 5.**
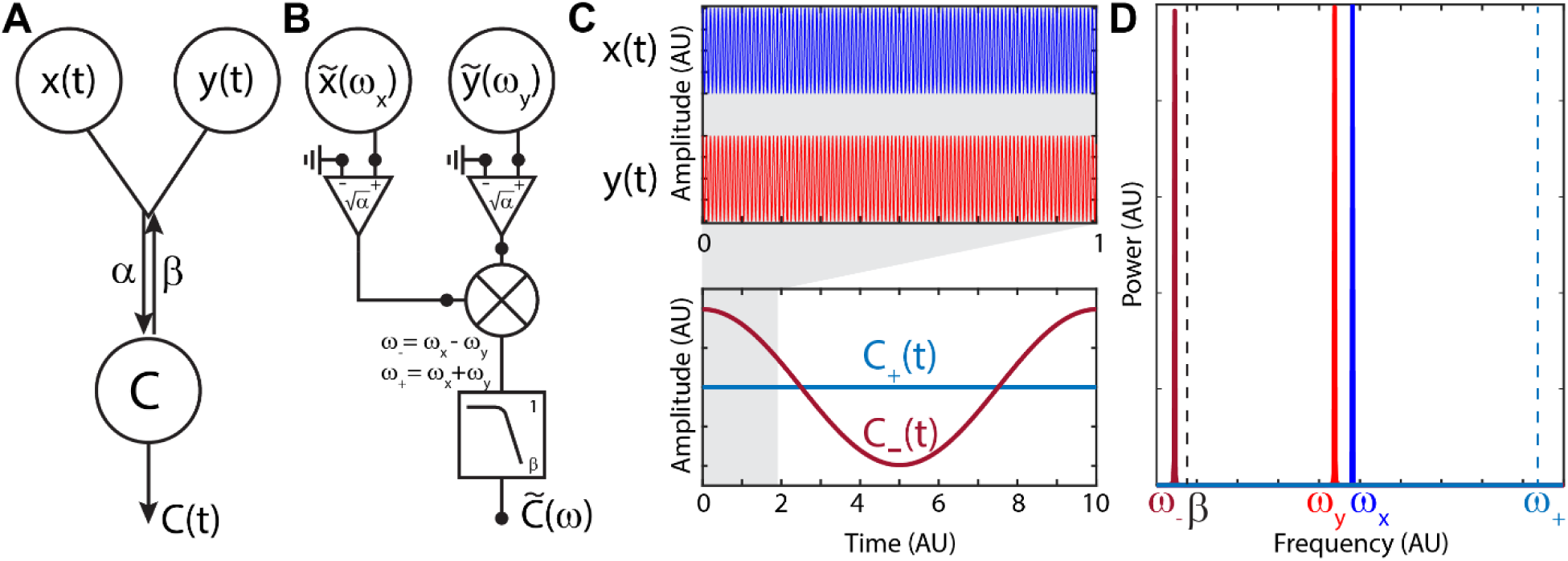
Multimerization as a nonlinear frequency mixer in series with low pass filter. (**A**) Two components, *x*(*t*) and *y*(*t*), that oscillate with frequencies *ω_x_* and *ω_y_*, dimerize with rate *α* to form a functional complex, *C*. *C* spontaneously disassembles with linear rate *β*. (**B**) The complex *C* acts as a frequency mixer (cross symbol) in series with a low pass filter. The spectral output of the complex includes frequencies that are all possible sums and differences of the input frequencies. These signals are then passed through a low pass filter whose frequency cutoff is set by the disassembly rate of the complex. (**C**) Top: Example time traces of *x* and *y*, *ω_x_* = 628.32 rad/t, *ω_y_* = 627.69 rad/t. Bottom: High [*C*_+_(*t*), blue] and low [*C*_+_(*t*), red] frequency components of the output signal *C*(*t*). Note that the x-axis scale on the bottom is 10x that of the top; *ω*_+_ = 1256 rad/t, *ω*_−_ = 0.63 rad/t, *β* = 1 t^−1^. (**D**) Power spectrum of (C). Blue and red: *ω_x_* and *ω_y_*, respectively; Dark blue and red: *ω*_+_ and *ω*_−_, respectively; Frequency cutoff, *β*: black dashed line. Location of *ω*_+_ indicated by dashed line, but power is negligible due to being above the frequency cutoff.

We apply the trigonometric identity

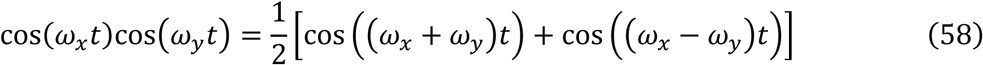

and integrate. The steady state response is

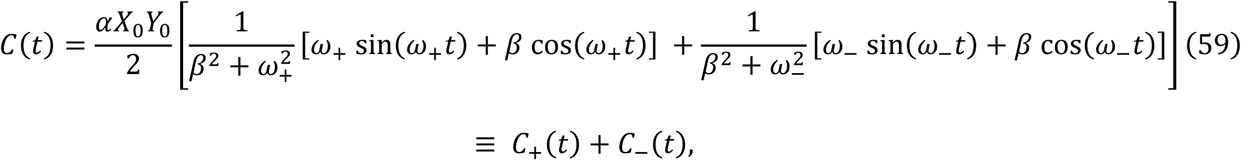

where *ω*_±_ = *ω*_1_ ± *ω*_2_. The output spectrum of the functional complex therefore contains frequencies that are the sum and difference of the input frequencies of the subunits. Moreover, each term is multiplied by a frequency dependent amplitude that functions as a low pass filter with frequency cutoff set by *β*, the disassembly rate of the complex (**FIGURE 5B**). If the original two oscillators are high frequency, *ω*_1_, *ω*_2_ ≫ *β*, the amplitude of the signal component with frequency *ω*_+_ ≫ *β*, *C*_+_(*t*), will be negligible. Furthermore, if *ω_x_* and *ω_y_* are very close in value, the beat frequency *ω*_−_ can be arbitrarily small such that *ω*_−_ ≪ *β*. This low frequency signal, *C*_−_(*t*), will pass the filter unimpeded (**FIGURE 5C,D**). Moreover, below the frequency cutoff the term in phase with the original oscillations will dominate, *i.e.*, the cosine term in **eq. 59**.

Generally, if a complex is assembled from multiple subunits, *a*, *b*, *c*, …, each with a Fourier spectrum composed of sets of angular frequencies, 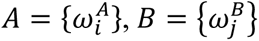, …, the above arguments can be applied iteratively to show that the output angular frequencies of the assembled complex will consist of all possible sums and differences of the input angular frequencies, 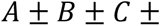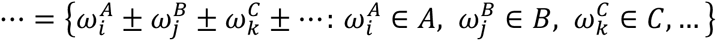 Any resulting frequencies below the cutoff set by the disassembly rate will appear in the complex’s spectrum.

For example, consider a homomultimeric complex consisting of *m* subunits of *x*, each oscillating at a single angular frequency *ω*_0_ with amplitude *X*_0_. We can consider the assembly of the complex as occurring either step-by-step one subunit at a time, or cooperatively with all subunits assembling simultaneously. First, and more simply, if the subunits assemble cooperatively with rate *α*, *i.e.*,

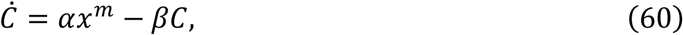

then the complex will generate a spectrum composed of a train of harmonics at even multiples of ω_0_.

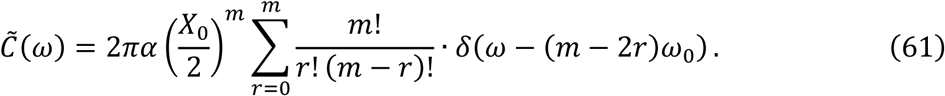

Conversely, if the complex assembles step by step, to simplify the problem we will assume that the individual subunits are numerous enough that complex formation and disassembly does not substantially affect their levels over time, such that *C*_1_ = *x*(*t*) = 〈*x*〉 + *X*_0_cos(*ω*_0_*t*). Then each subcomplex composed of *n* subunits obeys

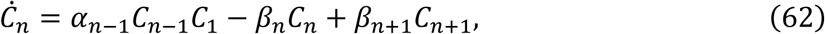

with *C_m_*_+1_ = 0. The first nonlinear term will again generate harmonics of *ω*_0_ by the same frequency mixing mechanism as above. However, because layers feed into each other from above and below by assembly and disassembly, all complexes will exhibit an infinite Fourier series of harmonics of *ω*_0_,

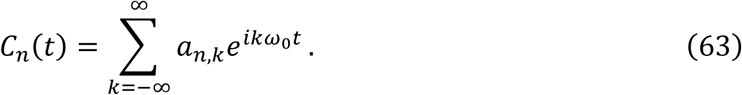

This can be treated by Floquet theory, but it is not possible to find a closed form finite analytic expression for the Fourier coefficients *a_n_*_,*k*_ that determine the amplitude of each harmonic. For example, with *m* = 2 this problem is mathematically identical to the equations of motion of ions confined in a Paul trap ^46^, where the solutions are expressed in terms of Mathieu functions that are notoriously unpleasant ^47^. For *m* > 2, the solutions are multidimensional generalizations of Mathieu functions and the behavior of the complex is most easily studied by numerical simulation.

Either assembly mechanism provides a simple way to generate low frequency harmonics such as those observed in p38 ^1^ that, moreover, is similar to the high degree of multimerization that occurs in formation of the myddosome complex in the earliest stages of interleukin and Toll-like signaling ^48–50^. Such harmonic frequencies are useful because they are orthogonal and can provide the minimal amount of crosstalk between signaling channels ^51, 52^.

### Generating and Transmitting Low Frequency Oscillations through Protein Cascades

Next, we demonstrate how combinations of these elements in complex, interconnected protein cascades can generate different frequency oscillatory signals from noisy input from the environment, and how the frequencies of those signals can be manipulated by the cascade to drive specific biological responses. This model is not meant to directly recapitulate the organization and behavior of the MAPK cascade or any other explicit network of interacting biological components. In any case, it is not clear that the dynamic behaviors of the individual components of such networks have been measured in sufficient detail to permit such an approach.

An example of such a network is shown in **Figure 6A**, and its equivalent representation in **Figure 6B**. The activated receptor activates two high frequency two-component oscillators, which then amplify signals at their resonant frequencies. These oscillators dimerize, combining the original two high frequency signals and generating and isolating a single low frequency, long period signal as a heterodyne. Next, these dimers oligomerize; oligomerization will generate harmonics of the low frequency signal from the heterodyne, any of which below the disassembly rate of the complex will survive. Thus, starting from a random input of white noise, stimulated complex assembly has resulted in generation of a long train of precise, well-defined harmonics as multiples of a low frequency fundamental.

**Fig. 6.**
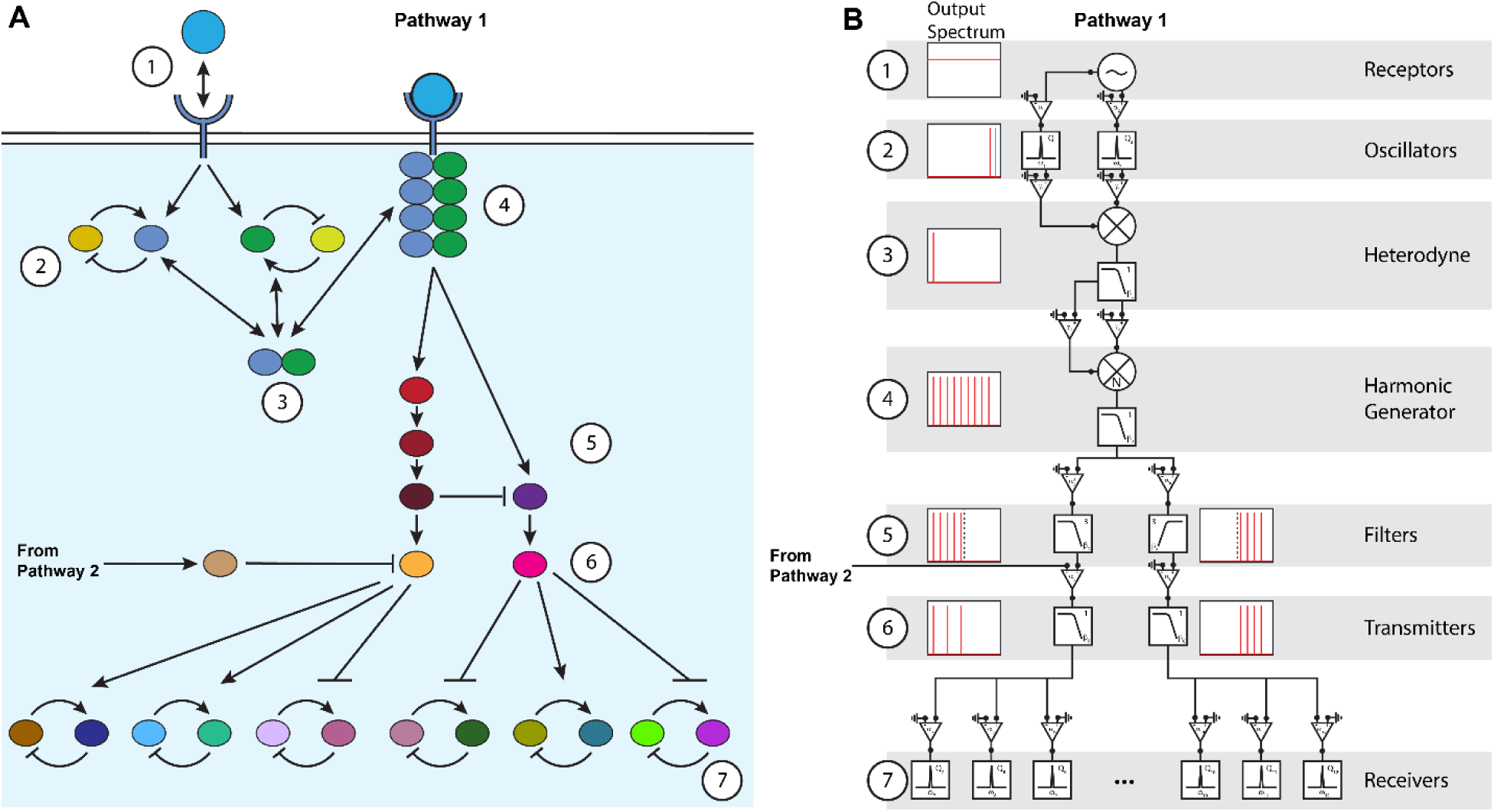
Generating and transmitting low frequency oscillations through protein cascades. (**A**) Biological and (**B**) equivalent representation of a complex, multilayered protein cascade that generates, sculpts, and transmits a spectrum of low frequency harmonic oscillatory signals to specifically control gene expression. (1) An environmental signal (*e.g.*, a cytokine) binds and unbinds at a receptor, generating a stochastic Poissonian white noise signal. (2) The bound receptor activates two high frequency oscillators acting as resonant amplifiers isolating two frequencies *ω*_1_ and *ω*_2_. (3) These oscillators dimerize, acting as a heterodyne to generate and isolate a low frequency signal *ω*_−_ = |*ω*_1_ − *ω*_2_|. (4) The dimers *N*-fold oligomerize at the activated receptor, generating *N* harmonics of *ω*_−_. (5) The harmonics are passed through layers of proteins acting as *m*^th^-order low and high pass filters to sharply divide high and low frequencies. (6) The low frequency signals are passed to one master regulator (*e.g.*, p38 ^1^), while the high frequency signals are passed to a second master regulator (*e.g.*, ERK1/2 ^28, 29^). An additional component from a separate Pathway 2 represses the low frequency master regulator to remove additional harmonics, fine-tuning the signal. (7) The master regulators act upon oscillating transcription factor receivers through biochemical resonance to activate specific genetic pathways ^1^.

Subsequent layers of the network can serve to further tune and distribute these harmonic signals. The spectrum is next passed through a few layers of proteins acting as an *m*^th^-order low pass filter and a high pass filter, to sharply separate and send low frequency harmonics to one master regulator, *e.g.*, p38 MAPK ^1, 24^, while high frequencies are sent to a second master regulator, *e.g.*, ERK1/2 ^19, 28, 29^. Finally, these master regulators select and activate specific transcription factor substrates in a signal frequency-dependent manner by biochemical resonance ^1^. At any stage, components from other signaling pathways can interact with elements to introduce, enhance, or suppress specific frequency signals to fine tune the spectrum (**FIGURE 6**).

For example, following the rules outlined above, the magnitude of the output spectrum of the low frequency master regulator is

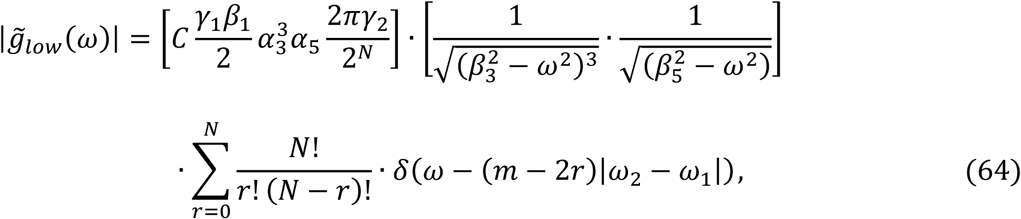

and that of the high frequency master regulator is

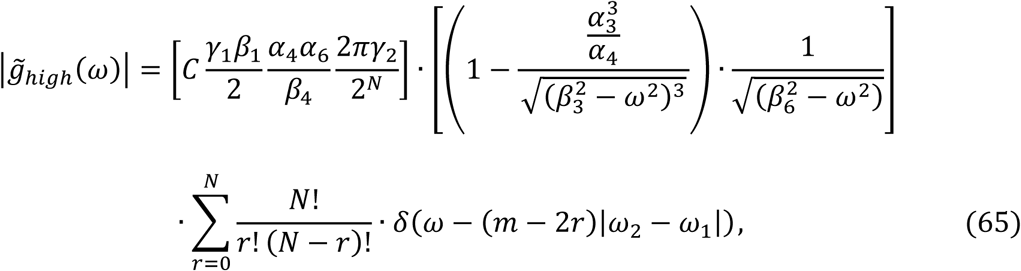

Here 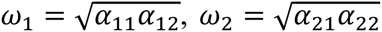 and *N* represents the number of subunits that make up a complex of oscillators assembled after stimulation. The first term in brackets represents a collection of coupling and degradation constants that sets the overall amplitude of the spectrum. The second term represents the modulation resulting from the filters separating the spectrum into high and low frequency components. Finally, the last term represents the collection of harmonics that result from combining the two high frequency oscillators activated by the receptor into a functional signaling complex.

## DISCUSSION

Here we have developed a graphical and mathematical language that provides both qualitative and quantitative descriptions of how different frequency signals can be acted upon by biological components. Some of these results are not novel. For example, it has long been appreciated that chemical reactions can act as filters ^39, 53^. However, the experimentally validated observation that some biological components are transmitting information encoded in different frequency oscillations ^1^ means that analogies to electronic telecommunications systems and information encoding and transmission as different frequency waves are more literal than previously appreciated.

Hence, it is advantageous to extend the analogies between biological and electronic network analysis as far as possible to provide a simple framework through which to understand the interactions of components in complex biological networks and cascades. To do this, we have here developed a simple graphical language, where each component and their interactions can be represented by standardized symbols. Each of these symbols corresponds to a quantitative expression of its impedance that characterizes its effect on a signal passing through it. Then, the mathematical expressions for the impedances of each element can be combined based upon how the components are connected in the network to quantitatively describe the overall effect of the entire cascade on the transmitted signal.

In developing this graphical and mathematical treatment, it has been our goal to keep the representations as simple as possible. It is our hope that the qualitative understanding provided by this simple graphical representation of complex biological networks might lead to its adoption and use even in contexts that do not require precise, quantitative levels of understanding. Recognizing that different collections of components result in behaviors that recapitulate standard elements in telecommunications systems, *e.g.*, filters, amplifiers, op-amps, and frequency mixers, allows one a qualitative understanding of how a complex network might behave without any quantitative analysis whatsoever. For example, the network we constructed in **FIGURE 6** was first developed by qualitative reasoning of how signals could be generated and how successive layers of the cascade could act upon those signals to roughly recapitulate our observations in p38 MAPK system ^1^. Only after we first laid out this qualitative roadmap did we then determine the quantitative parameters that yield the quantitative description.

## Acknowledgments

We thank Nigel Goldenfeld, Edward C. Cox, Paul Wiggins, Hernan Garcia, Matthew Scott, Prue Talbot, Nathan Gabor for useful discussions. This study was supported by startup funds provided by University of California, Riverside, the National Science Foundation (NSF POLS-2609970), and the UCR School of Medicine Dean’s Collaborative Seed Grant.

